# Activity-driven extracellular volume expansion drives vertebrate axis elongation

**DOI:** 10.1101/2022.06.27.497799

**Authors:** Arthur Michaut, Alessandro Mongera, Anupam Gupta, Mattia Serra, Pietro Rigoni, Jong Gwan Lee, Felipe Duarte, Adam R. Hall, L. Mahadevan, Karine Guevorkian, Olivier Pourquié

## Abstract

The vertebrate *bauplan* is primarily established via the formation of embryonic tissues in a head-to-tail progression. The biomechanics of this elongation, which requires the presomitic mesoderm (PSM), remains poorly understood. Here, we find that avian PSM explants can elongate autonomously when physically confined *in vitro*, producing a pushing force that can largely account for the posterior elongation of the embryo. Tissue elongation results from volumetric expansion that is driven by cellular activity and accompanied by inhomogeneous increase of the extracellular fraction along the AP axis. We show that FGF signaling promotes glycolysis-dependent production of Hyaluronic Acid (HA), which is required for expansion of the posterior PSM. Our findings link body axis elongation to tissue expansion through the metabolic control of extracellular matrix production downstream of FGF signaling.

**One-Sentence Summary:** Active tissue expansion propels body elongation independent of cell proliferation-driven growth

## Main Text

In the vertebrate embryo, the formation of sequential axial domains (head-neck-trunk-tail) requires the local addition of new material through cell proliferation, cell growth, and extracellular matrix (ECM) deposition, the rearrangement of existing material—i.e., cells, tissues, and ECM—or a combination of both (*1*). Classical experiments with frog explants have demonstrated the ability of anterior axial tissues to elongate autonomously along the anteroposterior (AP) axis (*2*). This behavior is based on convergence-extension (CE) driven by cell intercalation, a form of tissue rearrangement that is not observed in the avian embryo during elongation of posterior structures such as trunk and tail (*1, 3*). Posterior elongation ultimately relies on volumetric growth, with terminal addition of new cells from the neuromesodermal progenitor (NMP) zone located in the tail bud, the posterior-most region of the embryo (*1*). However, the force underlying this process does not depend on cell addition, as surgical ablation of the NMPs has negligible effects on elongation speed at timescales of the order of few hours. Similarly, impairment of cell proliferation both in chicken and zebrafish does not impact axial extension at short timescales (*4, 5*).

Several studies identified the posterior domain of the paraxial mesoderm, i.e., the presomitic mesoderm (PSM), as a major player in the process of posterior elongation (*4, 6*–*8*). The PSM appears as bilateral strips of mesenchyme that originate from the NMPs. In its anterior extremity, the PSM undergoes sequential segmentation into somites, epithelial blocks that contain the vertebral and skeletal muscle precursors (*9*). A posterior-to-anterior gradient of random cell motility is established in the PSM downstream of FGF signaling in all vertebrates analyzed (*4, 6, 8, 10*). In the chicken embryo, this motility gradient is hypothesized to control the generation of forces driving elongation of the posterior body (*4, 11*). The proposed mechanism is akin to pressure generation by confined gas particles (the motile cells) in response to an effective temperature gradient (FGF signaling). Due to the confinement of the PSM by tissues of higher cell density such as the lateral plate, the axial structures, the anterior PSM, the ectoderm and endoderm, the motility gradient is predicted to result in a posteriorly oriented pushing force (*4, 11*). Moreover, the pressure resulting from high motility in the posterior PSM is predicted to compress the neural tube and notochord, propelling the tail bud PSM precursors posteriorly and promoting the exit of new PSM cells, allowing to sustain elongation movements (*12*).

To investigate the central role of the PSM during axis elongation, we first probed the PSM ability to elongate autonomously when cultured *ex vivo* (Fig. 1, A to D). Dissected PSMs were transferred to non-adherent culture dishes and monitored for 3 hours. Their total length (from somite S1 to the end of the explant) rapidly decreased and the tissue adopted a typical pear-like shape, with the width of the posterior region, which is positive for the early PSM marker Mesogenin-1, increasing over time (Fig. 1, B to D). Unlike *ex vivo* cultures of *Xenopus* mesoderm (*2*), no autonomous CE is observed in these isolated PSMs. Therefore, maintenance of tissue shape as well as its elongation require surrounding tissues, either as a geometrical constraint or as a source of instructive, morphogenetic signals. These results further argue that tissue mechanical properties are not uniform along the AP axis, as previously reported for zebrafish, in which a gradient of viscosity and stiffness is observed along the PSM (*6, 8, 13, 14*).

**Fig. 1.**
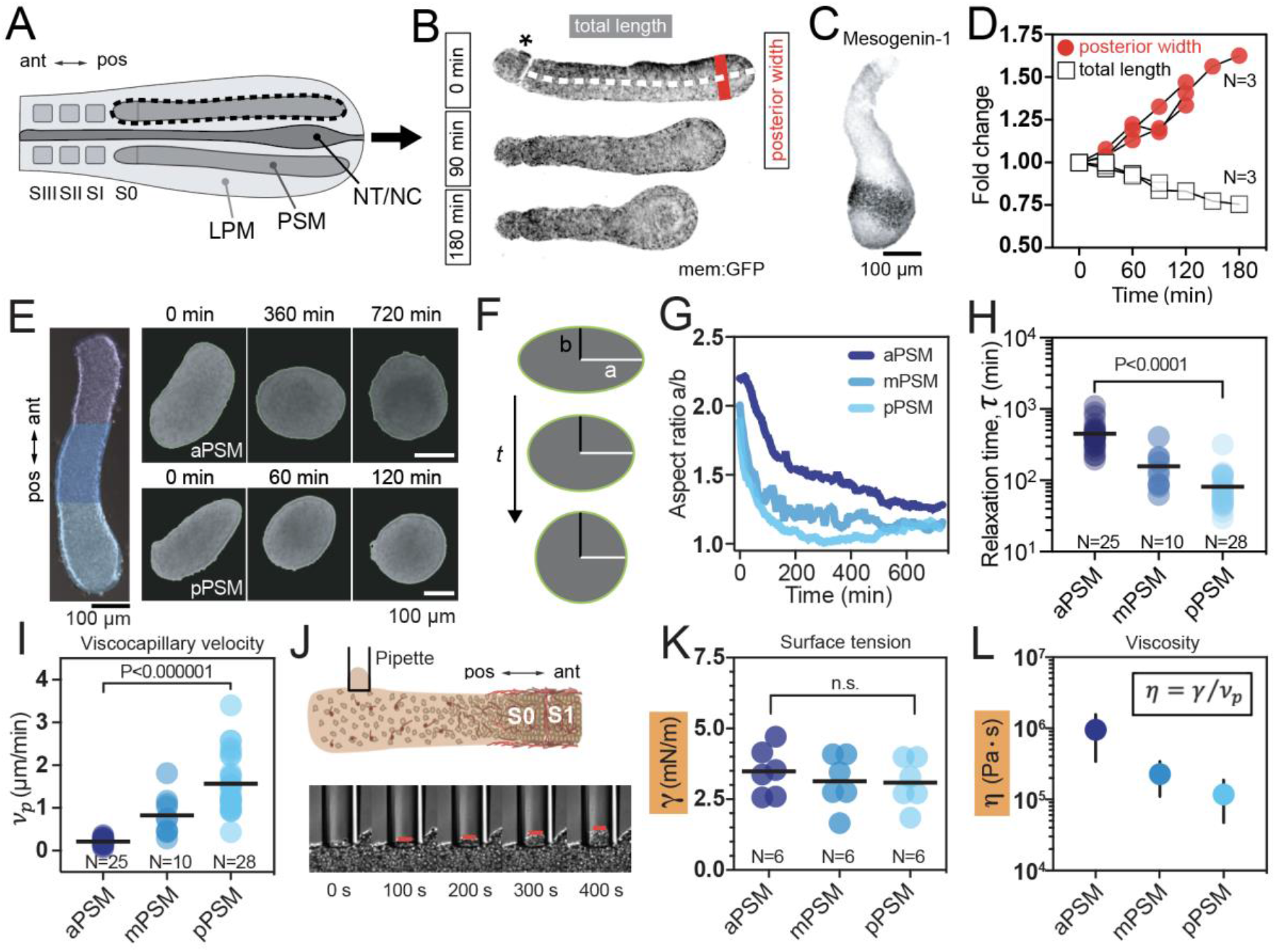
Fluid behavior of PSM explants. (**A** to **D**) PSM explants acquire a pear-like shape in culture. (A) Sketch of the anatomical features of the posterior body in chicken. LPM, lateral plate mesoderm; PSM, presomitic mesoderm; NT, neural tube; NC, notochord. S0-S3 indicate somite number. pos, posterior; ant, anterior. (B) Shape changes over 3 hours of a mem:GFP PSM explant in culture. Asterisk indicates somite SI. (C) Antibody staining for Mesogenenin-1 indicating that the expanding region has a posterior PSM identity. (D) Quantification of shape change, red dots indicate the posterior width (highlighted also in B) and white squares indicate total length (from somite SI to the end of the explant). (**E** to **I**) Quantification of viscocapillary velocity, *ν*_*p*_. (E) Examples of rounding of anterior (aPSM) and posterior (pPSM) explants. (F) Sketch showing how the aspect ratio changes during explant rounding. a, major semiaxis; b, minor semiaxis. (G) Plots showing examples of the evolution of the aspect ratio a/b for the different regions. (H) Gradient of relaxation timescale with higher values in the aPSM (bars indicate mean; P<0.0001, Mann-Whitney test). (I) Gradient of *ν*_*p*_ with significantly higher values in the pPSM compared to the aPSM (bars indicate mean; P<10^−6^, Mann-Whitney test). (**J** tand **K**) Measurement of surface tension, γ using the micropipette technique. (J) Sketch of pipette aspiration experiment with representative time series. (K) Plot of surface tension showing no significant difference between aPSM and pPSM (bars indicate mean; P=0.5606, Mann-Whitney test). (**L**) Viscosity (*η*) calculated from surface tension and *ν*_*p*_ (mean±SD).

To shed light on the mechanical parameters underlying PSM elongation *in vivo*, we set out to quantify spatial variations in tissue mechanical properties along the AP axis. We first quantified PSM fluid behavior by analyzing the rounding dynamics of explants from different regions (*15*– *17*) (Fig. 1, E and F). Rounding of the PSM explants is analogous to the behavior of a fluid parcel that tends to minimize its surface area/volume ratio to reach a final radius of *R*_*f*_ due to the influence of interfacial forces at its boundary (*15*). The evolution of the aspect ratio has a decaying exponential profile with a relaxation time given by 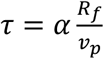, where *ν* _p_ is known as the visco-capillary velocity. *ν* _p_ corresponds to the ratio of surface tension over viscosity *γ/η and α* ≈ 0.95 when the tissue viscosity is much larger than the medium viscosity (*15*). The relaxation time is significantly larger for the anterior explants (aPSM) compared to the posterior explants (pPSM), varying from 453.8±213.7 to 81.75±54.37 min, P<0.0001 (Fig 1, F to H). Consequently, we obtain a posterior-to-anterior decreasing gradient in *ν* _p_ (from 1.57±0.67 to 0.22±0.08 μm/min, P<0.000001) (Fig. 1I).

To obtain absolute values of viscosity from the measured *ν*_p_ values, we performed micropipette aspiration experiments to quantify tissue surface tension (γ) *ex vivo* (*18*) (Fig. 1J and fig. S1) and found little variation of γ along the AP axis (from 3.09±0.84 for posterior to 3.5±0.85 mN/m for anterior, P=0.5606) (Fig. 1K). This therefore indicates that tissue-scale viscosity changes considerably during PSM differentiation, with a difference of almost 1 order of magnitude between the posterior (9.57×10^5^ ± 6.13×10^5^ Pa·s) and anterior PSM (1.18×10^5^ ± 0.7×10^5^ Pa·s) (Fig. 1L). Taken together, our results suggest that tissue viscosity largely accounts for the non-uniform mechanical response of the PSM to deconfinement.

To better understand the parameters giving rise to the fluid behavior in our explants, we turned to a minimal model of the underlying processes by building on a recent theoretical and computational description of body elongation driven by a combination of a gradient in cell motility and lateral confinement (*11*) (Fig. 2, A to D). To mimic the explant, we start with a fixed number of cells, but with an A-P gradient in cell activity (motility); the entire system is assumed to be confined by a soft rectangular boundary in two dimensions (Fig. 2A)(see SI for details).

**Fig. 2.**
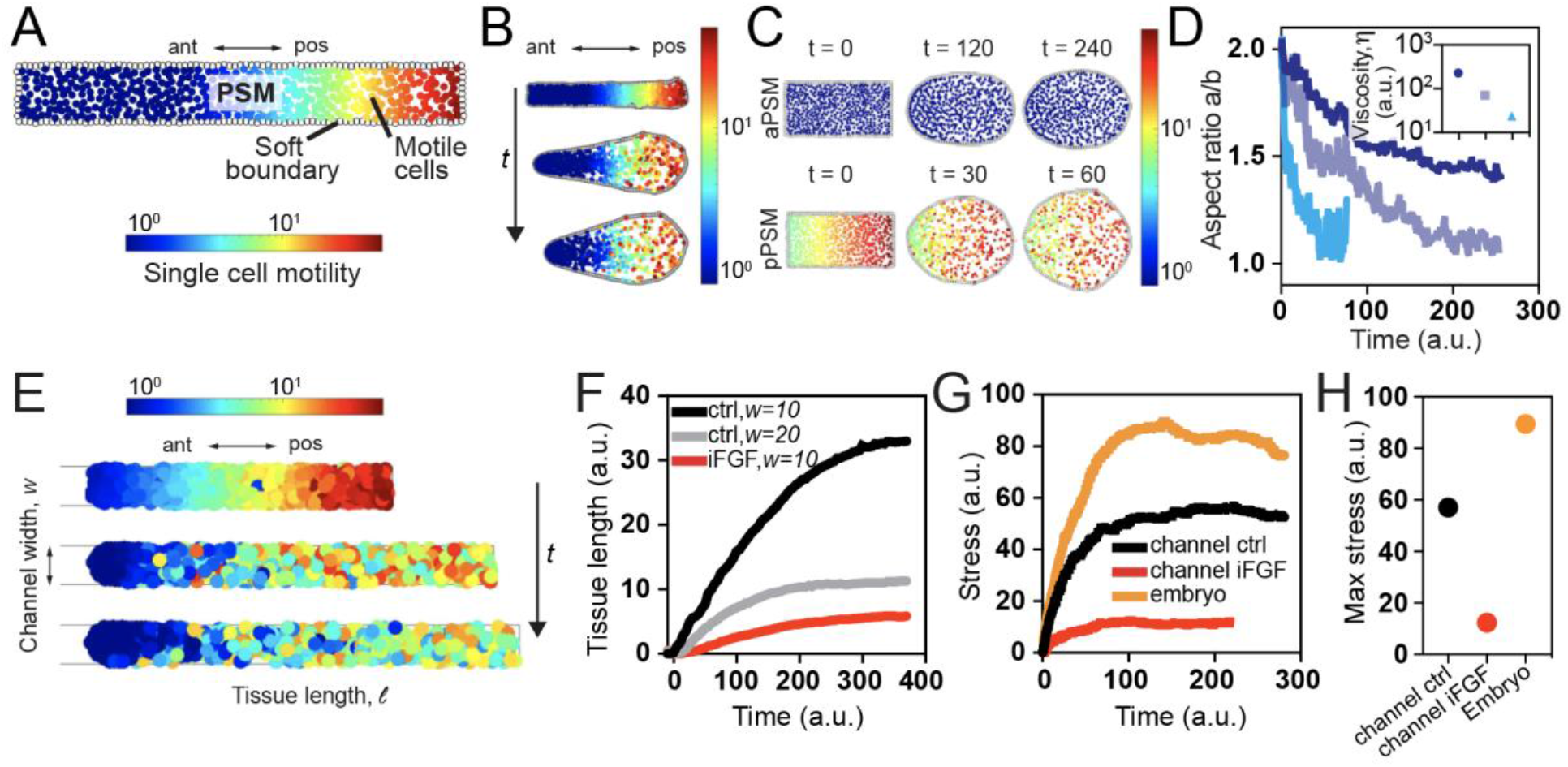
Computational model predicts PSM elongation upon confinement. (**A** to **D**) Computational model explains experimental observations. (A) Cells in the explant are modeled as colloidal particles surrounded by a deformable boundary. Cells display an AP gradient in activity (motility) and a mutually repulsive interaction. (B) Simulation of PSM explant in culture acquiring a pear-like shape due to an AP gradient in cell motility that leads to differential swelling of the boundary. This result is consistent with experimental observations described in Fig. 1B. (C) The tendency for cellular aggregates to round up is a function of their location along the AP axis as shown in Fig. 1E. (D) Simulations of the evolution of the aspect ratio a/b for explants from different regions. Inset shows the values of viscosity calculated from the simulations. (**E**) Confinement to a narrow and stiff channel leads to explants elongation in our simulation. (F) As the confinement is increased (from 20 to 10 cell size unit, black and grey line, respectively), the dynamics of elongation changes. Elongation in the channel is impaired when motility is reduced, mimicking FGF inhibition (red line). (G) Comparing the propensity for force generation by the elongating embryo (with a source of cells at the PSM posterior boundary, orange line) and the channelized explant (without a source of cells) with normal or reduced motility (black and red line, respectively). (D) Maximum elongation stress.

Our simulations show the emergence of a pear-shaped PSM upon deconfinement coupled to variations in cell density, consistent with experimental observations in Fig. 1B. We also simulated the rounding of cellular aggregates from posterior, mid and anterior PSM regions and found that posterior explants round up faster than those from the anterior region (Fig. 2, C and D), in agreement with our experimental results in Fig. 1E,F. We then extract the effective drift and diffusivity of cells (see SI) which, in combination with an active generalization of the fluctuation-dissipation relation (i.e., *D ∼ M/f*, with *D* = diffusivity, *M =* activity (motility), *f* = friction) allowed us to extract the effective viscosity of the cellular aggregates as a function of location along the AP axis (Fig. 2D, inset). The computational results are consistent with the experimentally inferred variations in the effective viscosity along the AP axis shown in Fig. 1I, confirming that gradients in activity (*M*), interpreted as variations in *f* are sufficient to explain the observed behavior. We note that in our model there are no variations in either the effective elasticity or the interfacial tension of our active colloidal aggregates along the AP axis (see SI for details), confirming that our minimal model suffices to explain our observations of the effect of the AP motility gradient.

We next asked whether PSM confinement by surrounding tissues, coupled to fluidity of its posterior region, can drive elongation. We first took advantage of our model and tested the role of confinement *in silico*. We replaced the soft confining boundary in our previous simulations with a stiff one characterized by a defined width, *w* (see SI for details). This simple change leads to tissue elongation in our simulations (Fig. 2E). The dynamics of this elongation depends on the channel width, with narrower channels resulting in longer explants and faster growth (Fig. 2F). The gradient in cell motility is required for this elongation, as reduction of FGF-dependent cellular activity leads to minimal growth (Fig. 2F). Furthermore, our model predicts that the elongational stress—that is, the pushing force produced by the explant during elongation— depends on cell motility and, to a much lesser extent, on addition of new cells (Fig. 2, G and H). This suggest that the force driving elongation is primarily produced by cell motility and gradients therein rather than new cells added from the NMPs.

To test the predictions of our simple model, and reconstitute the physical constraints of the embryo, we designed a PDMS microchannel array with channels of varying widths (75-160 μm) and heights (100-150 μm), allowing to impose different levels of mediolateral (ML) and dorsoventral (DV) confinement to PSM explants (Fig. 3A). With a 1.8 mm length, the channels are longer than PSM explants (500-1000 μm) and thus allow their confined elongation along the AP axis. PSM explants inserted in the channels elongate and form epithelial somites at their anterior extremity, suggesting that tissue differentiation proceeds normally in this artificial environment for up to 8 hours (Fig. 3B). The explants first exhibit a short contraction phase (characterized by a minimum length ℓ_min) until they reach full confinement in the channel. This is followed by an extension phase (Fig. 3C), during which the length of the whole explant (ℓ) increases. We analyzed the elongation dynamics by measuring the length increase after the contraction phase (ℓ − ℓ_*min*_) at high and low confinement, using channels of 90 μm and 160 μm widths with identical height (100 μm). In narrower channels, explants extend fast in the first 4 hours with speed 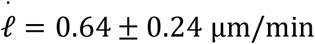 before they slow down (Fig. 3, C and D). In contrast, explants in wider channels grow linearly at a lower speed (0.15 ±0.08 μm/min, P=0.0286) (Fig. 3, C and D). Taken together, our results show that physical confinement is sufficient to promote elongation of the PSM *ex vivo* with elongation dynamics that depends on the degree of confinement. An important corollary of these findings is that molecular signals from adjacent tissues *in vivo* are not necessary for uniaxial elongation.

**Fig. 3.**
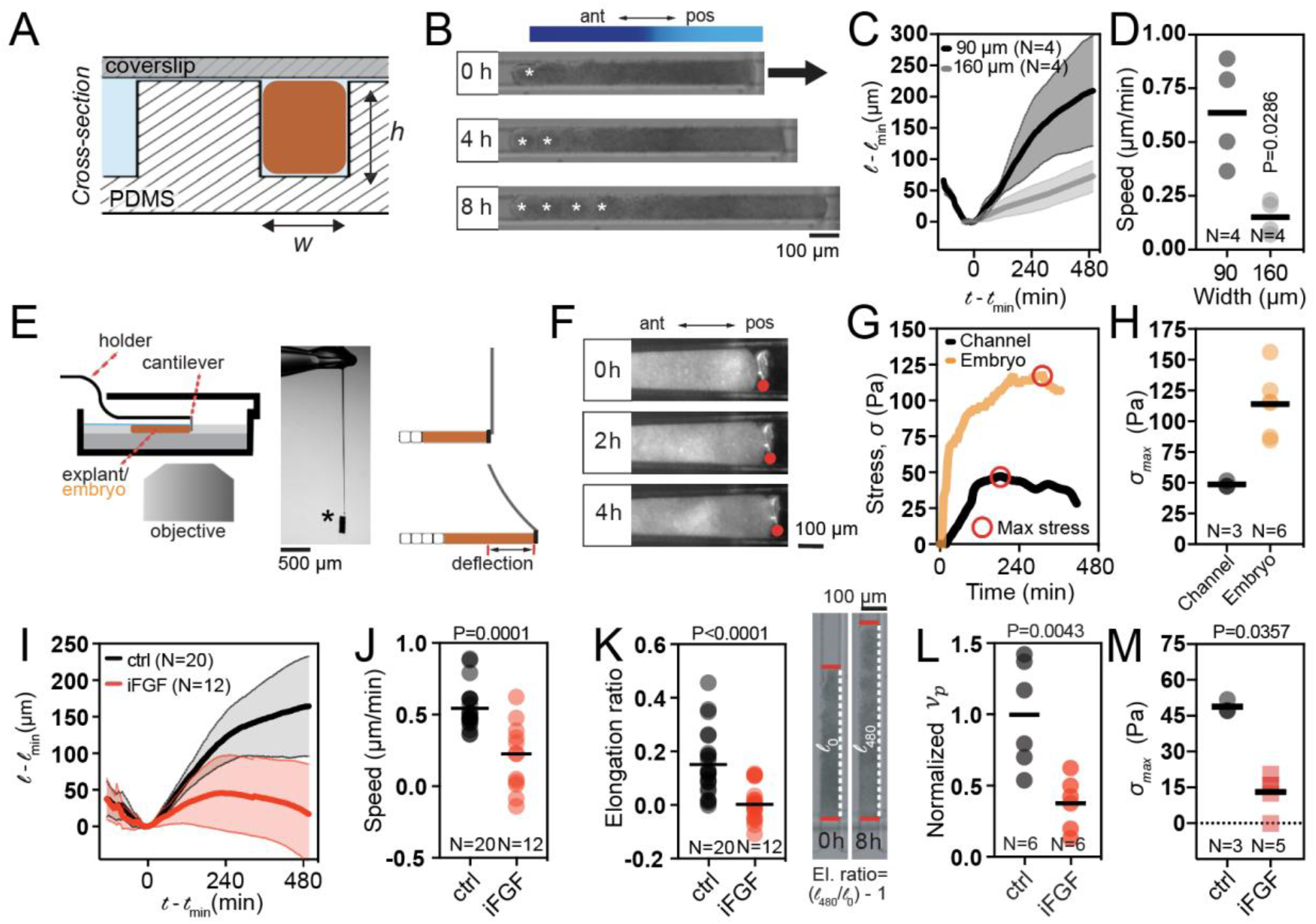
PSM explants elongate upon confinement and produce a force sufficient to drive axis elongation. (**E** to **H**) Experimental evidence of explant elongation upon physical confinement. (E) Sketch of the cross-section of a single channel with an inserted explant. (F) Example of PSM explant that elongates for 8 hours in the channel while forming somites in its anterior region. (G) Elongation dynamics of PSM explants in narrow (90 μm) and large (160 μm) channels. Mean and SD are plotted. (H) Elongation speed. (**I** to **L**) Measurement of stress produced by PSM explants *ex vivo* and by the whole embryo *in vivo*. (I) Sketch of the cantilever set up. Asterisk indicates the rigid aluminum force collector that is attached to a flexible glass capillary. The deflection of this glass is used to measure the stress exerted by the tissue. (J) Example of cantilever deflection in the channel. The position of the cantilever cross section is highlighted with a red dot. (K) Examples of cantilever deformations as a funtion of time in both explants (black) and embryos (yellow). The red circles indicate the maximal stress exerted during each experiment, (L) Plot of maximal stress produced by the explants and by the embryo. (**M** to **Q**) FGF signaling is required for proper elongation in the channel. (M) Dynamics of explant elongation upon FGF inhibition (iFGF) with PD03. (N) Elongation speed. (O) Elongation ratio. (P) Normalized viscocapillary velocity *ν*_*p*_. of posterior explants. (Q) Maximal stress produced. Mann-Whitney test.

While our results highlight the intrinsic ability of confined PSM explants to elongate, whether this can produce sufficient forces for posterior axis elongation as suggested by our simulations remains to be established. To address this issue, we devised a force sensor made of a rigid aluminum force collector attached to a flexible cantilever (Fig. 3E). We positioned the cantilever vertically at the opening of a microchannel of 140 μm width and 150 μm height. The cantilever was placed in contact with the posterior extremity of a growing explant and its deflection δ with respect to its initial position was monitored over time (Fig 3F).

We found that the maximum stress *σ*_*max*_ produced by a single confined explant during elongation is 49 ± 3 Pa (Fig. 3, G and H). To compare the force produced by the PSM *ex vivo* to that generated by the whole embryo, we inserted the cantilever vertically in the region posterior to the axial progenitors in embryos cultured *ex ovo*. We found that the whole embryo, including medial tissues and the two bilaterally symmetrical PSM columns, produces a stress of 114 ± 26 Pa (Fig. 3, G and H). Thus, the derived force generated by the PSM could account for a significant part of the posteriorly-directed force generated by the whole embryo.

These measured stresses are in good agreement with predictions of our simulations (Fig. 2, F to H) and the continuum model presented previously (*11*).

We next investigated the role of FGF/MAPK signaling on *ex vivo* elongation, fluidity, and force generation (Fig. 3, I to M). When PSM explants were cultured with the MEK inhibitor PD0325901 (PD03) which inhibits FGF signaling in the PSM, the average elongation speed decreased dramatically from 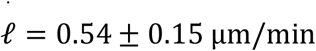 to 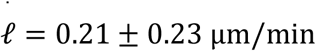 (P=0.0001)(Fig. 3, I and J). To quantify the effect of PD03, we calculated the elongation ratio, 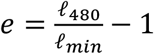, which describes how much the explant has elongated after 8 hours (480min) compared to its initial length ℓ_*min*_. While control explants are characterized by a 15% increase of their length (*e* = 0.15 ± 0.12), treated explants show minimal elongation (*e* = 0.01 ± 0.06) (Fig. 3K). We also assessed whether MAPK inhibition impacts tissue fluidity and the force generated by the explants *in vitro*. Upon treatment with PD03, we found a ∼60% reduction in *ν*_p_, indicating that FGF signaling is required to maintain high fluidity within the posterior PSM (Fig 3L). We also detected a 75% reduction in the maximal stress exerted by the explants (*σ*_*max*_ = 13 ± 8 Pa, P=0.0357) (Fig. 3M). Taken together, our results show that the PSM is the major source of force production during posterior body morphogenesis and that this process requires active FGF/MAPK signaling.

In vivo, the high-FGF posterior region of the PSM was shown to be a major elongation driver. We thus asked whether the anterior and posterior regions of the PSM (aPSM and pPSM) contribute differentially to its elongation *ex vivo*. We set out to dissect PSM explants of different lengths, reducing progressively the amount of aPSM included (Fig. 4A). We reasoned that the elongation ratio *e* should be correlated with the explant length if the PSM is formed by an active and a less active region. We found that shorter explants, where the fraction of pPSM *vs*. aPSM is larger, display higher values of elongation ratio (Fig. 4A), suggesting that elongation is primarily driven by the pPSM.

**Fig. 4.**
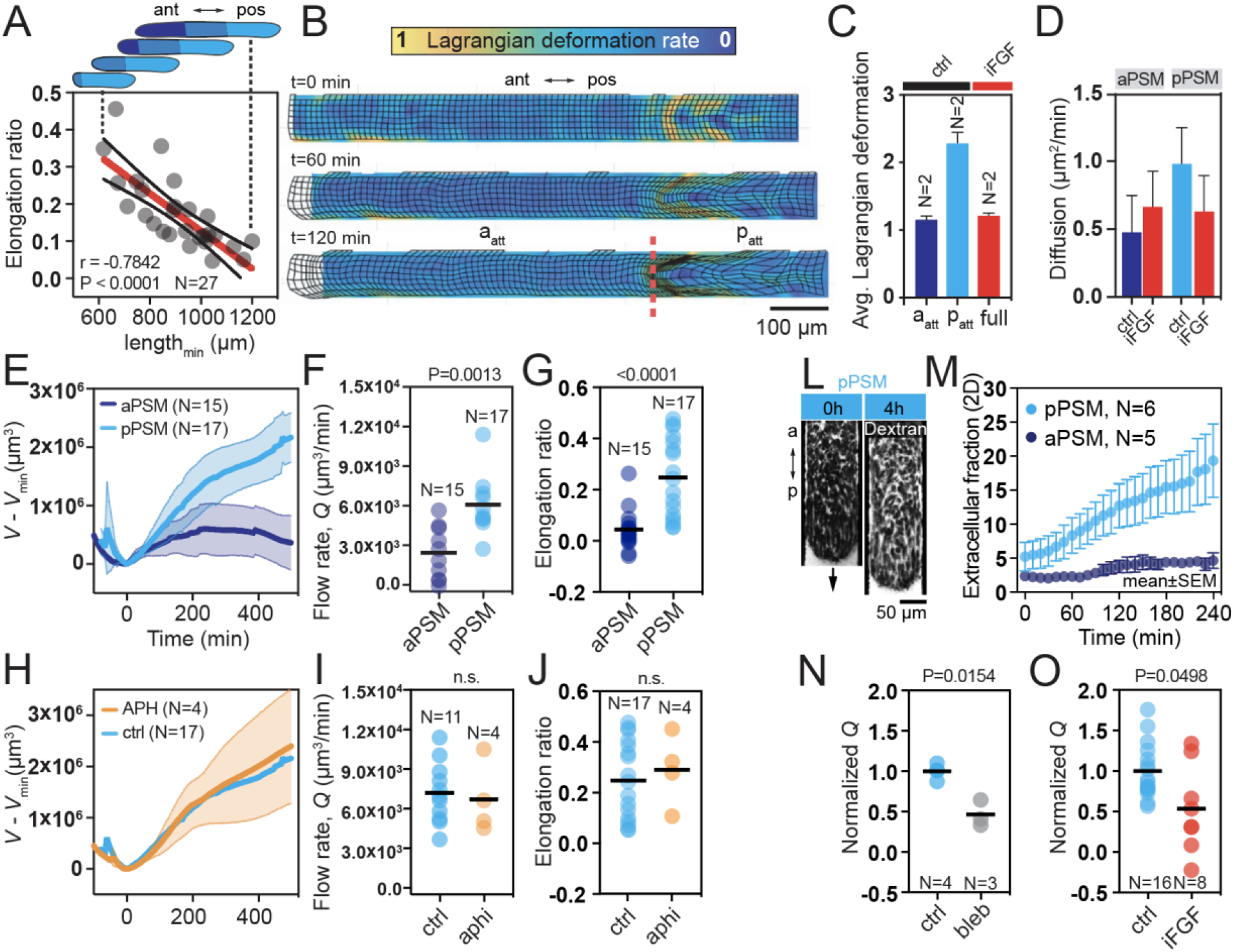
PSM explants elongate via volumetric growth dependent on cell motility. (**A**) Elongation ratio is negatively correlated with explant length. (**B**) Dynamic morphoskeleton (DS) analysis showing larger deformations in the posterior PSM during elongation in the channel. The dashed red line indicates the location of an attractor, which defines the boundary between an undeformed anterior region a_att_ and a posterior region p_att_ characterized by large isotropic and anisotropic deformation. (**C**) Plot of average Lagrangian deformation in control and PD03 treated explants. (**D**) Cell diffusion analysis measured by individual cell tracking. Right image shows a snapshot of cell trajectories. (**E** to **G**) Volumetric expansion of pPSM explants. (E) Dynamic of volumetric growth in aPSM and pPSM explants. (F) Plot of volumetric flow rate. (G) Plot of elongation ratio. (**H** to **J**) Volumetric expansion does not depend on cell proliferation. (H) Aphidicolin-treated (APH) pPSM explants show similar dynamics of volumetric growth. (I) Volumetric flow rate in treated pPSM explants. (J) Elongation ratio in treated pPSM explants. (**L** and **M**) Extracellular fraction (ECF) increases during pPSM elongation. (L) Example of pPSM elongation with labeled ECF with fluorescent dextran. (M) Plot showing ECF increase in the pPSM and no increase in the aPSM. (**N**) Normalized flow rate upon treatment with blebbistatin. (**O**) Normalized flow rate upon treatment with PD03.

To quantify the contribution of the pPSM in elongation, we used a Lagrangian deformation metric called dynamic morphoskeleton approach (DM) to investigate tissue-scale local deformations from Particle-Image-Velocimetry (PIV) data (*19*). These Lagrangian deformations are quantified by the backward finite-time Lyapunov exponent field (FTLE) and visualized through a deforming grid (see Methods and Fig. 4B). The DM on whole PSM identified large deformations— both isotropic- and shear-type deformations—in the explants’ posterior end, but not in the anterior region (Fig. 4B). A dynamic attractor, i.e., a moving region toward which the tissue converges, divides the posterior quarter of the PSM from the anterior region (Fig. 4B), representing an intrinsic boundary between PSM domains that undergo contrasting levels of Lagrangian deformations. By computing the spatial average of the maximum Lagrangian deformation field 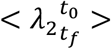, we found that the pPSM deforms on average twice as much as the aPSM, which remains undeformed (Fig. 4C and Methods). We also compared cell motility in the two regions and found larger diffusivity in the posterior region, in line with the gradient of motility observed *in vivo* (*4, 6*) (from 0.48 ± 0.27 μm^2^/min to 0.98 ± 0.27 μm^2^/min)(Fig. 4D). Analysis of explants upon MAPK inhibition revealed reduced deformation in the posterior region, resulting in uniform 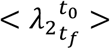 along the entire explant (Fig. 4C). Blocking FGF activity results also in a 36% reduction of cell diffusivity in the posterior PSM (Fig. 4D). Taken together, these data show that the pPSM region, which exhibits high cell diffusion and low viscosity, is characterized also by large deformations during elongation. Moreover, both motility and deformation depend on active FGF/MAPK signaling. The presence of isotropic deformation is in line with previous 2D analysis *in vivo* which identified a region of the posterior PSM that is subjected to in-plane expansion (*20*).

To test whether this expansion in 2D corresponds to a volumetric growth, we analyzed the elongation dynamics of isolated anterior and posterior domains cultured in microchannels (Fig. 4, F to J). By analyzing our data in 3D, we found that posterior explants increase their volume in the AP direction during the entire experiment, while volume expansion rapidly stops in anterior explants (Fig. 4, E to G). We found that the volumetric flow rate (*Q*), which corresponds to the amount of material passing through a given cross-sectional area per unit time, is more than 2-fold higher in the pPSM compared to the aPSM (from 2426 ± 2005 to 6094 ± 2288 μm^3^/min)(Fig. 4F). In addition, elongation ratios are significantly different (from 0.04 ± 0.08 to 0.24 ± 0.15, P<0.0001)(Fig. 4G). These results argue that the PSM elongates by volumetric growth in its posterior region and ruled out the potential role of CE movements during elongation.

We next investigated the role of cell proliferation in the control of PSM expansion. Previous analyses *in vivo* showed no posterior bias in cell proliferation rate (*20*) and no significant reduction of elongation speed upon chemical inhibition of cell division (*4*). We did not observe any significant difference in pPSM elongation, including elongation ratio (P=0.6353) and volumetric flow rate (P = 0.7567), upon inhibition of cell division with aphidicolin (Fig. 4, H to J).

An alternate possibility is that the volume increase of the pPSM is caused by dilation of the tissue in response to high cell motility, as observed for a gas when temperature increases. To test this hypothesis, we first quantified the extracellular fraction (ECF) by injecting fluorescently-labeled dextran in pPSM explants (Fig. 4L). After a 4-hour culture period, we observed an almost 4-fold increase of the ECF in pPSM explants (from 5 to 20%) (Fig. 4, L and M). Thus, the volume increase of the pPSM mostly corresponds to an increase in extracellular space. Next, we impaired the motility of PSM cells by adding blebbistatin to the culture medium as previously described (*4, 6*). We observed a 50% decrease in the volumetric flow rate in treated pPSM explants (Fig. 4N). A similar impairment of volumetric flow rate was observed upon FGF inhibition (Fig. 4O). Taken together, our results suggest that high cell motility downstream of FGF signaling results in an ECF increase in pPSM explants driving their elongation in a confined environment.

*In vivo*, we observed a posterior to anterior gradient of ECF, with larger ECF in the pPSM (Fig. 5, A to C). The extracellular environment of the mesenchymal pPSM cells is composed of water, ions and extracellular matrix (ECM), the composition and mechanical properties of which are under tight spatiotemporal regulation. As cell motility in 3D requires a substrate for traction/friction (*21*), we reasoned that the ECM might also contribute to growth. However, the ECM can serve not only as a passive substrate for cell motility but also as an active expandable filler. One of the most abundant ECM components is Hyaluronic Acid (HA), a negatively charged polymer that forms a viscoelastic gel promoting hydration and lubrication in adult tissues (*22*). By swelling with water, HA can generate compressive osmotic forces that have been implicated in the morphogenesis of embryonic tissues (*23, 24*). The Hyaluronan synthase *Has2* mRNA is highly expressed in the most posterior and anterior regions of the PSM, as well as in the somites (Fig. 5D). We next showed that HA concentration is uniform along the AP axis using an ELISA immunoassay (Fig. 4E). We performed solid-state nanopore analysis of the molecular weight distribution of HA polymer chains (Fig. 5, F and G). This analysis revealed a shift towards larger HA molecules in the pPSM (1100±228 kDa) compared to the aPSM (739±221 kDa), suggesting a capacity to achieve higher degrees of swelling in the former (*25*).

**Fig. 5.**
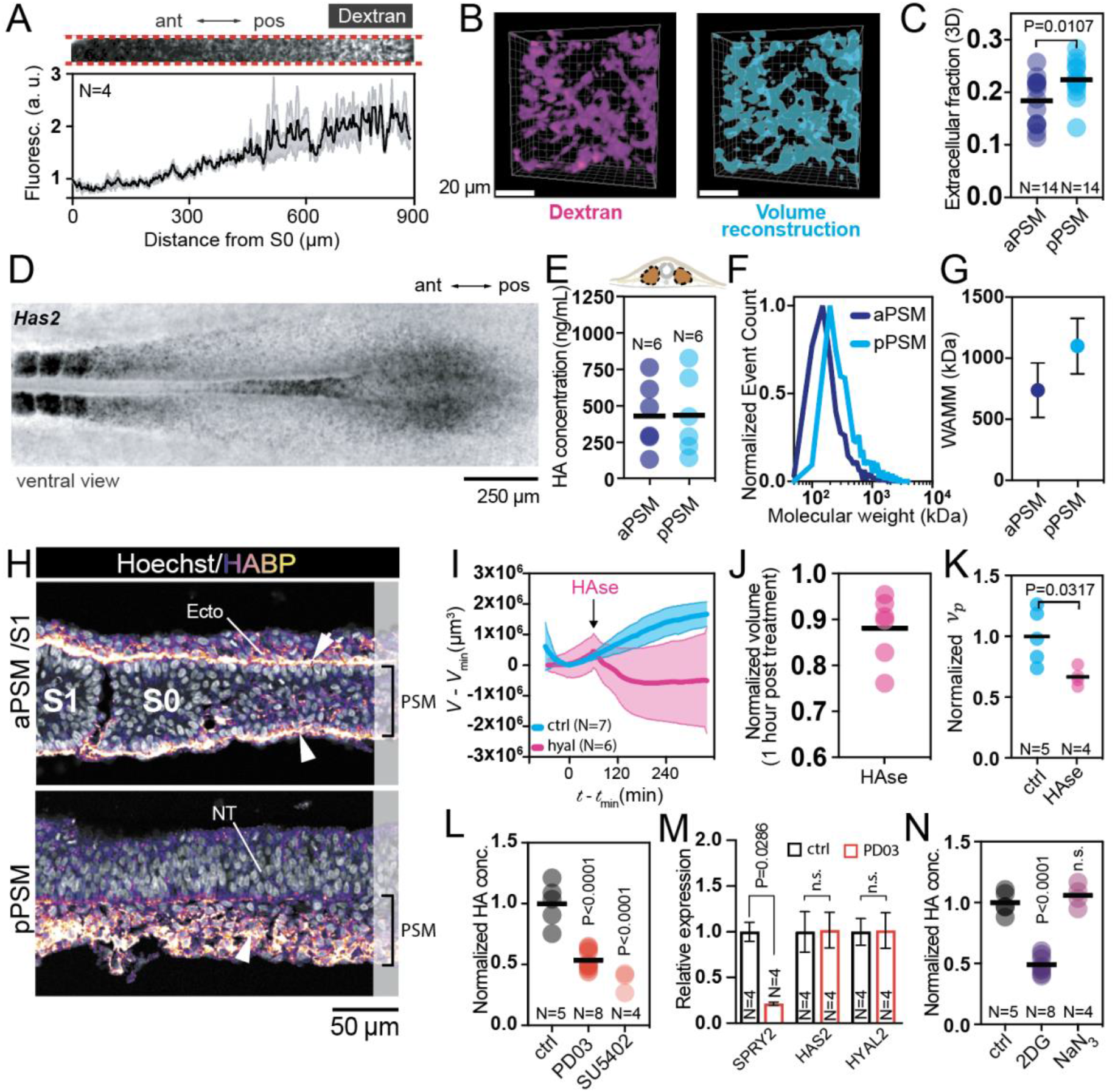
PSM expansion requires glycolysis-dependent HA synthesis downstream of FGF signaling. (**A** to **C**) A posterior-to-anterior gradient of ECF *in vivo*. (A) Plot of dextran signal intensity along the PSM *in vivo*. (B) 3D rendering of dextran signal and its volume recontruction. (C) Plot of 3D ECF *in vivo*. (**D**) HCR in situ for Has2. (**E**) ELISA assay for HA concentration along the PSM. (**F** and **G**) Quantification of HA polymer length via solid phase nanopore analysis. (F) Plot showing normalized event count for polymers of different lengths. (G) Plot of weighted average molecular mass (WAMM). (**H**) Sagittal sections of staining with a biotinylated HA-binding protein (HABP). (**I** to **K**) Effects of hyaluronidase treatment on explant growth and fluidity. (I) HA degradation blocks volumetric growth. (J) 12% volume reduction in explants treated for 1 hour with HAse. (K) Plot of normalized *vp* upon HAse treatment. (**L** to **N**) FGF signaling controls HA production via glycolysis. (L) Normalized HA concentration in the PSM upon treatment with PD03 or SU5402. (M) qPCR analysis of Has2 and Hyal2 upon PD03 treatment. (N) Normalized HA concentration in the PSM upon treatment with 2DG or sodium azide.

We next analyzed HA localization using a biotinylated HA-binding protein (HABP). While in the pPSM, HA is present only in the bulk of the tissue, in the aPSM it is primarily located at its surface (Fig. 5H), similar to what has been reported for fibronectin (*26*). To test the role of HA, we treated PSM explants in the microchannels with 300 U/mL hyaluronidase, an enzyme that degrades HA. HA degradation blocked volumetric growth and resulted in a 12% volume reduction after 1 hour (Fig. 5, I and J). Moreover, we found that HA degradation leads to a significant decrease in *ν*_*p*_ of the posterior explants, suggesting that HA may contribute to lubricate the PSM and maintain high fluidity of its posterior region (Fig. 5K). Thus, HA is required for the maintenance of PSM architecture and its volumetric expansion.

FGF signaling has been shown to control HA production in different developmental contexts, primarily through upregulation of Hyaluronic acid synthase 2 (*Has2*)(*27*–*29*). To test the role of FGF signaling in the regulation of HA synthesis, we treated 10-12 somite embryos for 5 hours with either PD03 or the FGF receptor inhibitor SU5402 and assessed HA concentration by ELISA immunoassay on dissected pPSM. Upon inhibition of FGF signaling, we detected a 50-65% reduction in HA concentration (Fig. 5L). We next tested whether FGF signaling controls HA levels by regulating the expression of genes coding for enzymes involved in HA production/degradation, such as *Has2* and *Hyal2*, respectively. Quantitative PCR analysis revealed no difference in the expression levels of both genes upon FGF inhibition (Fig. 5M), indicating that FGF signaling does not control HA levels by transcriptionally activating core enzymes involved in HA regulation. HA synthesis requires the production of UDP-sugars via glycolysis, a metabolic pathway that has been shown to sustain body axis elongation downstream of FGF signaling (*30, 31*). This glycolytic activity also regulates cell motility in the PSM (*30*).

We therefore tested whether FGF may affect HA levels through regulation of aerobic glycolysis. Upon inhibition of glycolysis with the hexokinase inhibitor 2DG we found a 50% reduction in HA levels (Fig. 5N), indicating that an intact glycolytic flux is required for HA production during posterior axis elongation. Importantly, no impact on HA concentration was found upon blockage of oxidative phosphorylation with NaN_3_. This argues for an unexpected link between FGF signaling, glycolysis and HA production in the regulation of tissue volumetric expansion and the force that enables posteriorly directed body elongation.

Together our observations along with a minimal computational model demonstrate that the posterior PSM acts as an autonomous force-generating engine driving elongation of the chicken embryo trunk. A gradient in FGF signaling promotes cell diffusion and glycolysis-dependent HA synthesis, which are both required for ECF expansion. This process results in volumetric growth of the tissue independently of cell proliferation. Whether HA serves as a substrate to physically support cell crawling or acts as a signal to promote ERK/MAPK-dependent cell motility as shown in cancer is still unclear (*32*). This FGF- and HA-dependent process of volume expansion might be at play during elongation of other mesenchymal structures such as the limb bud (*33*). Our work demonstrates how the crosstalk between cell signaling and metabolism can control passive and active tissue mechanics and force generation during morphogenesis of the embryo.

## Supporting information

Supplemetal data_model

## Acknowledgments

K.G. and A.M. would like to thank Françoise Brochard-Wyart for fruitful discussions. We would like to thank Aditi Chakrabarti, who helped with the early version of the cantilever experiments. This work was partially supported by the French Agence National pour la Recherche under grants No. ANR-14-CE32-0009-01 to K.G. and National Institutes of Health grants 5R01HD097068-04 (O.P., L.M.) and R01 GM134226 (A.R.H.) as well as the NSF Simons Center for Mathematical and Statistical Analysis of Biology at Harvard University (L.M.), the Simons Foundation (L.M.) and the Henri Seydoux Fund (L.M.).

## Author contributions

Conceptualization: AM, AM, LM, MS, KG, OP

Methodology: AM, AM, AG, JGL, PR, FRD, ARH, KG, LM, OP

Investigation: AM, AM, AG, LM, MS, KG, OP

Funding acquisition: OP, LM, KG

Project administration: OP

Supervision: OP, KG, LM

Writing – original draft: OP, AM, AM, LM

Writing – review & editing: OP, AM, AM, KG, LM

## Data and materials availability

All data, code, and materials used in the analysis must be available in some form to any researcher for purposes of reproducing or extending the analysis. Include a note explaining any restrictions on materials, such as materials transfer agreements (MTAs). Note accession numbers to any data relating to the paper and deposited in a public database; include a brief description of the data set or model with the number. If all data are in the paper and supplementary materials, include the sentence “All data are available in the main text or the supplementary materials.”

## Supplementary Materials

Description of computational model.

## Materials and Methods

### Embryo preparation and dissection

Wild-type eggs were incubated for 40 hours at 37 **°**C, 65% humidity and embryos were harvested at stage HH11 (12-14 somites). We performed dissection in PBS with ***Ca***^**2+**^, ***Mg***^**2+**^ cations and collagenase type IV (Thermo Fisher Scientific diluted in PBS at 1 mg/mL in PBS) was used for 10 minutes. Dissection was carried out by means of pulled borosilicate rods (1 mm of diameter, Sutter instrument) pulled using a P97 filament-based pipette puller (Sutter instrument) and severed to yield roughly a 10 μm tip.

### Embryo culture

For regular time-lapse experiments, 10-somite stage embryos were harvested using the filter paper technique described by (Chapman et al. 2001) and cultured ventral side up. For force measurement experiments, we added a layer of mineral oil (Sigma) covering the embryo and the gel-based medium to keep the humidity and prevent the embryo from drying up as the cantilever was inserted. To facilitate elongation measurements, DiI staining for local labelling of landmarks was carried out by injecting a solution of DiI (Vybrant Multicolor cell-labeling kit, Invitrogen) by means of a glass needle.

### Tissue culture and treatments

The culture medium for explant culture consisted of DMEM-F12 (Life Technologies) supplemented with 10% FBS (VWR) and 1% penicillin-streptomycin (2mM L-Glutamine, 100U Penicillin, 100 g/ml Streptomycin). (-)-Blebbistatin (Sigma) stock solution was prepared by resuspending it in DMSO at 25 mM. It was then diluted in culture medium at 20 μM. PD0325901 (AXON MEDCHEM BV) stock solution was prepared by resuspending it in DMSO at 10 mM. It was then diluted in culture medium at 2 μM or 10 μM.

### Rounding experiments

Explants were cultured on LabTek dishes coated with 0.3 mg/ml PEG-PLL (PolyEthyleneGlycol-PolyLysin, PLL(20)-g[3.5]-PEG(2), SuSoS) in HEPES 10 mM for 1 hour. LabTek dishes were rinsed twice with PBS and incubated with culture medium at 37 °C and 7.5% *CO*_2_ for one hour prior to adding explants.

### Microchannel experiments

A PDMS microchannel array (see microchannel fabrication below) was placed in a 60 mm Petri dish. Microchannels were closed by a glass coverslip. Next, they were saturated with a PBS solution and gas was removed using a vacuum chamber. Microchannels were then coated with PEG-PLL by vigorously pipetting a 0.3 mg/ml PEG-PLL solution to replace water. After one hour of coating, the PEG-PLL solution was washed away by PBS. The microchannel array was then incubated with culture medium at 37 and 7.5% *CO*_2_ for one hour prior to adding explants. Explants were carefully slid into the channel by means of a fine glass rod. For the PSM force measurement in a microchannel, *CO*_2_ could not be controlled because of the cantilever insertion. The culture medium was therefore supplemented with 10 mM HEPES.

### Microchannel fabrication

An array of microchannels ∼1mm long, and with rectangular cross-sections varying between 75μm-170μm in width, and uniform height (either 100μm or 150μm) was fabricated in polydimethylsiloxane (PDMS) using a soft-lithography procedure. Because the channels were going to be used with the PDMS side on the coverslip (open grooves), care was taken to have the PDMS layer be less than 3mm for better optical resolution. Once the PSM explants were positioned inside the channels, the array was closed with a coverslip treated with 0.3mg/ml PEG-PLL (SuSos) solution to prevent cell adhesion.

### Cantilever fabrication and calibration for *in vivo* measurements

Rectangular probes of 800 μm by 400 μm were cut out of aluminum foil (25 μm thick) using a scalpel blade. The cantilever arm was made from glass capillary that was pulled using a laser puller to produce flexible tips of approximately 1 μm in diameter. The probe was fixed to the tip using a UV polymerizing glue (Bondic). The calibration was done by measuring the resonant frequency of the cantilever. Briefly, the cantilever was excited by a gentle blow and its oscillating response was recorded using a fast camera (Phantom v 7.3 at 8,000 frames per second) and extracted using the kymograph function in Fiji along a line perpendicular to the capillary. The resonance frequency was then obtained from the peak of the the power spectrum computed by Fourier transform, using the Python package Scipy (Jones et al. 2001). The stiffness of the cantilever, *k*, is related to the resonance frequency through 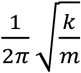 where *m* is the mass of the probe. We could fabricate cantilevers ranging from 1-100 mN/m.

### Cantilever fabrication and calibration for *in vitro* measurements

Force measurement in channels were carried out using a similar setup to the in vivo system. The glass cantilever was replaced by an Atomic Force Microscopy (AFM) cantilever (Bruker MLCT-O10 with spring constants ranging from 0.01-0.2N/m). Rectangular aluminum probes were fixed to AFM cantilever using a UV polymerizing glue (Bondic). We used the factory spring constant for the system stiffness.

### Deflection analysis

Cantilever deflection was measured by tracking the position of the cantilever tip over time using the ‘Manual Tracking’ plugin from the Fiji software. Deflection was defined as the distance of the cantilever tip with respect to its initial position.

### Stress measurement

We measured the force *f* produced by the elongation of the whole axis (in vivo) or individual PSM explants (in vitro) by tracking the deflection *δ* of our calibrated cantilever of stiffness *k*. We then calculated the force through *f* = *kδ*. To obtain the stress, which is force per area, *σ* = *f/A*, we approximated the area by the area of contact *A* = *w* × h with *w* the width of the probe and *h* the height of the probe in contact with the tissue. *In vivo*, we measured *h* by measuring the size of the insertion depth through the distance between the focal plane before and after insertion. Typically, *h* = 70 − 90*μm*, which is coherent with the known thickness of the embryo. *In vitro*, we directly used the thickness of the channel for *h*.

### Cell tracking

Motility in mmicrochannels was analyzed automatically tracking the PSM cells by means of the Fijji ‘TrackMate’ plugin. Cell trajectories were visualized using the Track Analyzer pipeline (Michaut 2022, in preparation).

### Aspiration setup

We use a setup that has been previously described in detail(*18, 34*). Briefly, pipettes were prepared by pulling 15 cm long borosilicate capillaries (outer diameter: 1 mm, inner diameter: 0.58 mm, Sutter Instruments), using a filament-based pipette puller (P-97 Sutter Instruments). We used pulling parameters to yield pipettes with almost parallel walls (parameters: 1 cycle P=500, HEAT=495, PULL=0, VEL=40, TIME=30). The tip of the pipettes were manually severed with a ceramic tile (Sutter Instruments) to obtain diameters ranging between 35-55 μm and fire-polished using a microforge (MF-1, TPI). Pipettes were subsequently coated with a 0.3 mg/mL PEG-PLL in HEPES solution (10 mM) for 1 hour and rinsed with double-distilled water. Aspirations were conducted with a hydrostatically controlled pressure system. A pipette was connected to a vertically displaceable water reservoir by means of a pipette holder (IM-H1, Narishige). The pipette holder was manipulated with a 3D micromanipulator (assembled using Velmex and Newport elements). The reservoir was displaced along a vertical rail (Newport and Thorlabs).

### Surface tension measurement

We measured the critical pressure *ΔP*_*c*_ above which an aspirated PSM explant starts flowing into the pipette. This critical point is defined when the aspirated length *L* > *R*_*p*_ with *R*_*p*_ being the pipette radius (Guevorkian et al 2010). As the geometry of the explant is not a sphere, we rewrote the relation between surface tension *γ* and *ΔP*_*c*_ using the explant local curvature *k* : 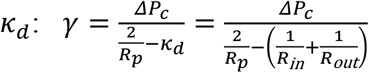 (see Fig. S1).

### Electroporation

Chicken embryos were electroporated *in ovo* at stage HH5. Eggs were windowed and a pCAGG-H2B-RFP plasmid solution (described by (Denans, Iimura, and Pourquié 2015)) at 1 μg final concentration and fast green (FCF 0.4, Sigma) was microinjected into the space between the vitelline membrane and the epiblast. Electroporations were carried out by placing a cathode below the embryo in the yolk and fine tip anode on the primitive streak at 70-80% of its total length away from its posterior end. Five successive squared pulses of 12V for 50 ms were produced using a BTX ECM 830 electroporator. Eggs were then taped to seal the opening and re-incubated for 24 hours.

### In vivo and ex vivo wide-field video-microscopy

Rounding and microchannels experiments were carried out on a computer controlled, wide-field inverted microscope (Zeiss Axioobserver Z1) equipped with a motorized stage and EMCCD camera (Evolve Photometrics). Explants were imaged by stitching together images (Zen 2 software stitching function) acquired with a 10X objective (EC Plan-Neofluar, numerical aperture=0.3). The acquisition rate was 10 frames per hour (6 min between frames). Temperature and *CO*_2_ were maintained respectively at 37 and 7.5% by means of a Pecon TempController 2000-2 and a Pecon *CO*_2_Controller 2000 (except for the PSM force measurement in a microchannel for which only the temperature was controlled).

For elongation experiments, embryos were cultured in a custom 6-well (one per embryo) sealed chamber and imaged on a microscope using a custom-built time-lapse station (Bénazéraf et al. 2010). We used a computer controlled, wide-field epifluorescent microscope (Leica DMR) workstation, equipped with a motorized stage and cooled digital camera (QImaging Retiga 1300i). For each embryo, several images corresponding to different focal planes and different fields were captured at every single time-point. The acquisition rate was 10 frames per hour (6 min between frames). For force measurement experiments, a single embryo was cultured and imaged on a computer controlled, wide-field inverted microscope (Zeiss Axioobserver Z1) equipped with an EMCCD camera (Evolve Photometrics). At each frame, one image was acquired with a 2.5X objective (N-Achroplan, numerical aperture=0.07) and one image with a 10X objective (EC Plan-Neofluar, numerical aperture=0.3). The acquisition rate used was 10 frames per hour (6 min between frames). When fluorescence was used (DiI, or H2B-RFP electroporated PSM), an HXP 120 C light source with a filter cube (excitation 565/30, emission 620/60) were used.

### Confocal microscopy

PSM explants were imaged by confocal microscopy on a Zeiss LSM780 confocal microscope using a 20X Plan-ACHROMAT (numerical aperture: 0.8) objective. A Z-stack was acquired with 4 μm interval between consecutive sections.

### Immunohistochemistry

Explants were rinsed with PBS and fixed in fresh PBS, 4% PFA for 2 hours at room temperature. Then, they were rinsed in PBS and incubated for an hour in PBS, 0.1% Triton. Explants were rinsed three times in PBS and incubated in blocking solution (PBS, 0.1% Triton, horse serum 1%) for 45 minutes. Explants were then incubated overnight at 4 with rabbit polyclonal anti-CMESPO antibody (Oginuma et al. 2017) (1:1000). Explants were then washed three times for 1 hour at room temperature in PBS, 0.1% Triton. Explants were blocked and secondary antibodies coupled with Alexa fluorophores (Life technologies) were incubated overnight at 4. Nuclear staining was performed with Hoechst33342 (1:1000, Life technologies). F-actin staining was performed with Phalloidin-Alexa 647 (1:50, Invitrogen). All immunostaining data were acquired using a Zeiss LSM780 confocal microscope using a 20X objective.

### Elisa assay

Embryos were first treated *ex ovo* for 5 hours with 50 μM PD0325901 (AXON MEDCHEM BV), 500 μM SU5402 (ApexBio), 2 mM 2-Deoxy-D-glucose (Sigma-Aldrich), and 1 mM Sodium azide, NaN_3_ (Sigma-Aldrich). 4-6 PSM were dissected from embryos placed in PBS (see ‘Embryo preparation and dissection ‘section) and pulled in a 1.5 ml Eppendorf tube with PBS on ice. PBS was removed and 50 μl of cell lysis buffer (Cell Lysis Buffer 2, R&D Systems, # 895347) were added. Tubes were incubated at 37 degrees for 15 minutes and tissue lysis was then facilitated by pipetting up and down the explants for 10 seconds. This operation was repeated 2 more times and was followed by a final incubation for 30 minutes. The protein concentration of each sample was then measured using a GENESYS 10S UV-Vis spectrophotometer (Thermo Fisher Scientific). Samples were diluted at 2 μg/ml using Calibrator Diluent RD5-18 (R&D Systems). Hyaluronan quantification was then performed using the Hyaluronan Quantikine™ ELISA kit from R&D Systems (DHYAL0).

### HA size distribution analysis

aPSM and pPSM specimens were separately diced and incubated with proteinase K (10μL at 20mg/mL U/mL, New England Biolabs, Ipswich, MA) at 37 °C - overnight to digest protein components, including structural elements and HA-binding proteins. An equal volume of a phenol:chloroform:isoamyl alcohol (25:24:1 v/v/v, Fisher Scientific, Hampton, NH) was added to the sample, mixed thoroughly, and centrifuged for 15 min at 14,000×*g* in a Phase Lock Gel Tube (QuantaBio, Beverly, MA) to separate HA and other aqueous components from the organic phase. The process was repeated once using pure chloroform to remove residual phenol.

For isolation, 150 μl of streptavidin magnetic beads (10 mg/mL Dynabeads M-280, Invitrogen, Carlsbad, CA) were washed three times with 1X PBS 0.05% Tween, washed three times with 1X PBS only, and then resuspended in 50 μl of 1X PBS. 41 μL of biotinylated versican G1 domain (bVG1, 0.363 μg/μl, Echelon Biosciences, Salt Lake City, UT) were then added to the beads, mixed, and incubated for 1 hour at room temperature on a rocker. After incubation, the beads were washed three times with 150 μl 1X PBS to remove unbound bVG1.

The bVG1-streptavidin beads were subsequently reconstituted with the solvent-extracted HA solution and incubated at room temperature for 1 hr with gentle rocking. HA-bound beads were pulled down by a magnet and washed three times with 1X PBS, after which 50 μl of measurement buffer (6M LiCl, 10 mM Tris-HCl, 1 mM EDTA, pH 8.0) was added directly to the sample to disrupt the VG1 binding pocket and release HA. Beads were again pulled down by magnet and the purified HA was removed and stored at -20°C until measurement.

For HA size analysis (*35*) a single nanopore device (7.9 nm, 20 nm thick membrane) obtained commercially (Norcada, Inc. Alberta, Canada) was rinsed with DI water and ethanol, dried with filtered air, and treated with air plasma (30 W, Harrick Plasma, Ithaca, NY) for 2 min on each side before being loaded into a custom 3D printed (Carbon, Redwood City, CA) flow cell and immediately interfaced with clean measurement buffer. Ag/AgCl electrodes (Sigma-Aldrich, St. Louis, MO) in each chamber were used for voltage application and ionic current measurement through a patch-clamp amplifier (Axopatch 200B, Molecular Devices, Sunnyvale, CA) which was used to check low-noise baseline current and confirm pore diameter. Purified HA in measurement buffer was then introduced to the *cis*-side of the membrane and a 300 mV bias was applied to induce molecular translocations. Trans-membrane ionic current was recorded at a rate of 200 kHz using a 100 kHz four-pole Bessel filter with an additional 5 kHz low-pass filter applied in post processing of data. Signals caused by translocations of HA through the pore into the opposite (*trans-*) chamber were defined as transient reductions in the ionic current >5X the baseline standard deviation and with durations in the range of 25 μs−2.5 ms. Molecular weights were assigned to each molecular translocation signal from its area using a calibration curve produced by measuring quasi-monovalent HAs (*36*) through the same kind of nanopore device.

### Dynamic Morphoskeletons

Given a planar velocity field **v**(**x**, t), we identify the Dynamic Morphoskeleton(*18*) (DM), and specifically attractors, from the backward Finite-Time Lyapunov Exponents (FTLE). We compute the FTLE as

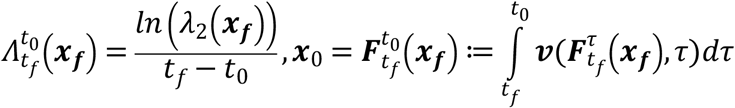

where *λ*_2_(***x***_***f***_) denotes the highest singular value of the Jacobian of the flow map 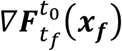 and 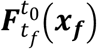 the flow map describing the trajectories from their final ***x***_***f***_ to initial ***x***_0_ positions. To compute the FTLE, we first calculate 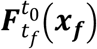 by integrating the cell velocity field **v**(x, t) using the MATLAB built-in Runge-Kutta solver ODE45 with absolute and relative tolerances of 10−6, linear interpolation in space and time, and a uniform dense grid of points ***x***_***f***_at the final time *t*_*f*_. Then we compute 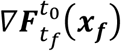 by finite differencing 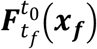 with respect to ***x***_***f***_ Finally, we compute *λ*_2_(***x***_***f***_) using the in-built MATLAB singular values function (see (*18*) for details). We note that 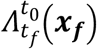 has units of 1/time hence describing deformation rates, while *λ*_2_(***x***_***f***_) is dimensionless and quantifies the maximum Lagrangian deformation of a region whose final position is ***x***_***f***_ during the time interval ***t***_0_, ***t***_*f*_. As aggregate deformation measure(*18*), we use the spatial average of 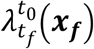, which can be computed as

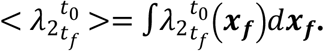

**Fig. S1.**
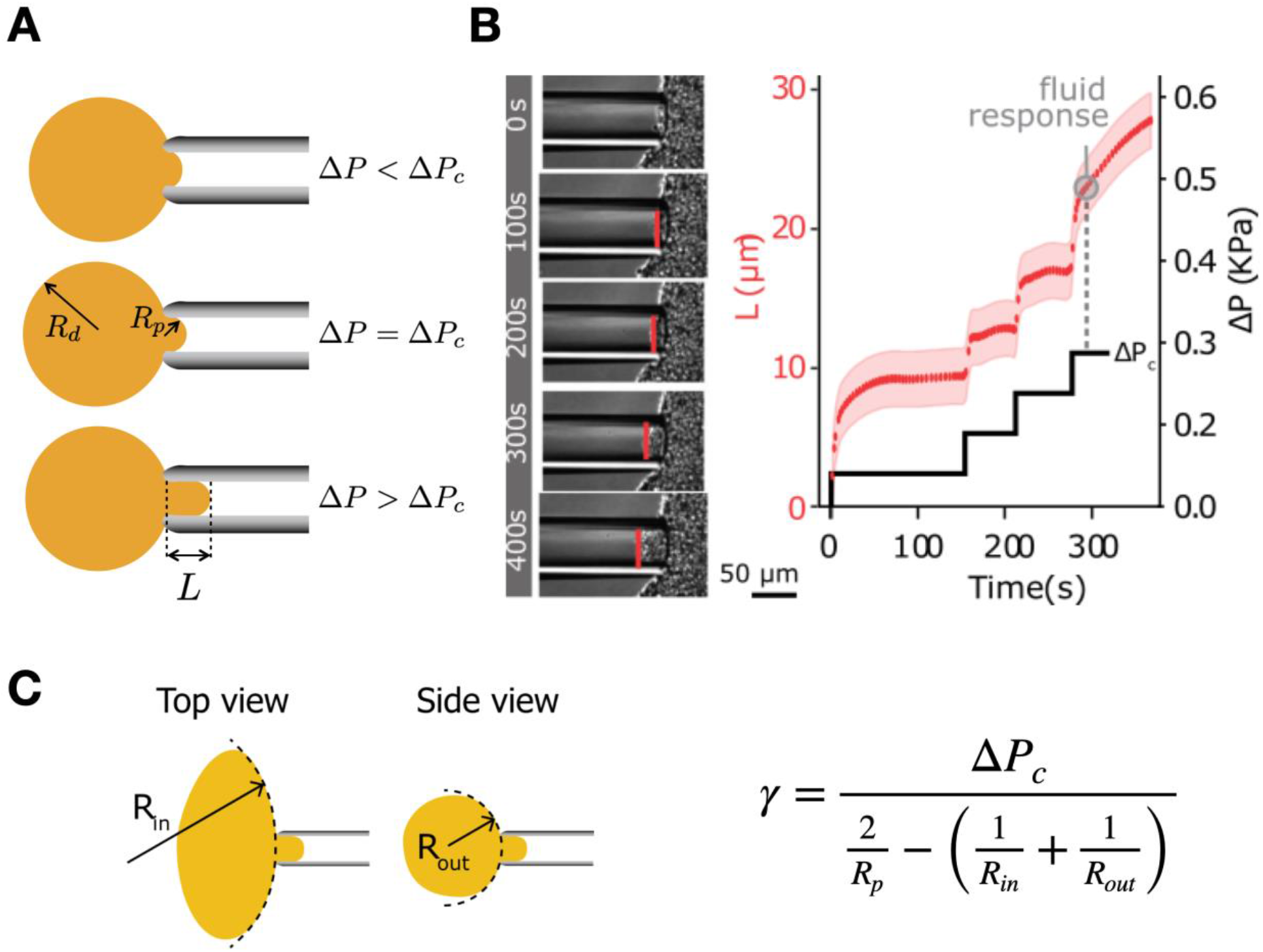
Surface tension measurement by pipette aspiration. (**A**) Definition of the critical aspiration pressure *ΔP*_*c*_, below which the curvature of the aspirated tissue increases up to a maximal value 2/*R*_*p*_ following the Laplace law. Above *ΔP*_*c*_, the curvature is maximal and the tissue flows into the pipette. (**B**) Example of measurement of *ΔP*_*c*_ by a sequence of steps of aspiration pressure *ΔP*, until the tissue flows. (**C**) Relation between a surface tension and *ΔP*_*c*_ for a non spherical tissue.

