## Supplementary material for "Activity-driven extracellular volume expansion drives vertebrate axis elongation": Supplemetal data_model

**Computational Model:**

To understand how the conditions for vertebrate embryo elongation emerge, we consider a minimal theoretical model of the system (Figure 2) starting from a system of active cells growing inside a confining passive soft boundary or a rigid channel walls. The active cells of the embryo are modeled using interacting soft circular discs of size $a$ subject to forces with the appropriate Langevin dynamics. Cells are assumed to be active with random motility analogous to a Brownian particle, but this movement of the cell is not related to the temperature of the environment but is instead due to the active nature of the cell [Berthier & Kurchan, 2013, Mallory et al 2014, Regev et al, 2022]. The cells interact with each other through an interaction potential such that they have short-range repulsion to avoid the overlap and mid-range attraction (two-cell size) and no long-range interaction (more than two-cell size) [Wang et. al. 2020]. The interaction potential between the cell is given by:

$$U(\boldsymbol{x}) = \frac{1}{2}\Sigma_j \Sigma_{i\neq j} u_{ij},$$

$$u_{ij} = \begin{cases} \epsilon\left(\left(\frac{a}{r_{ij}}\right)^2 - 1\right)\left(\left(\frac{r_c}{r_{ij}}\right)^2 - 1\right)^2 & \text{for } r_{ij} \leq r_c \\ 0 & \text{for } r_{ij} > r_c \end{cases}$$

where $r_c = 2a$; and $r_{ij}$ is the separation between the cells $i$ and $j$. $\epsilon$ is the depth of the potential well. The other contribution to the dynamics of the cell will come from the interaction from the boundary. The time steps to evolve the dynamics are chosen in such a way that the cells cannot escape these boundaries. So, if we combine the above three contributions, the equation of motion for a cell with coordinate $r_i$ can be written as:

$$\gamma\, \dot{\boldsymbol{r}}_i = -\frac{\partial U}{\partial \boldsymbol{r}_i} + \boldsymbol{\xi}_i(t) + \boldsymbol{f}_{ext}$$

where $\gamma$ is the tissue viscous friction, $\boldsymbol{f}_{ext}$ is the force which is exerted on the cell by the boundary, and $\boldsymbol{\xi}(t)$ is random force with zero mean and a variance related to its activity, i.e. $< \xi_i(t) >= 0$; $< \xi_{i,\alpha}(t)\xi_{i,\beta}(t') > = 2\,\mathcal{M}\mu_{\mathrm{t}}\delta(t-t')\delta_{\alpha\beta}$. Where $\mathcal{M}$ is the single cell motility (activity) and $\xi_{i,\alpha}$ are the $x$ and $y$ components of $\xi_i$. The viscous friction is a result of the interaction of cells with the extra-cellular matrix (ECM). Note that in our simulations the collisions are elastic but overdamped. We can use the results from statistical physics [Kardar 2007] and can relate the microscopic diffusivity of a (Brownian) cell to the activity by the relation $\mathcal{D} = \frac{\mathcal{M}}{\gamma}$. Analysis of cell motility in the chicken embryo presomitic mesoderm (PSM) [Bénazéraf et al., 2010] shows that there is an anterior-posterior gradient in the activity of cells. To incorporate this gradual reduction in the activity of the cells owing to reduction in FGF concentration anteriorly, we turn off the activity with a probability which changes exponentially with time;

$$\Pr = 1 - e^{-t/\tau}$$

and with space away from the tailbud along AP direction;

$$\Pr = 1 - e^{-x/L}$$

Where $\tau$ is the slowest time scale associated with the decay motility of the cell over time, and $L$ is corresponding length scale over which the motility of the cell decays. This length scale $L$ is directly related to the motility decay time scale through the relation $L \sim a\rho_0\ \mathcal{D}\ \tau$ [Regev et al 2022], where $\rho_0$ is the maximum packing fraction of the cells in the PSM. This model is extension of our work where we have used this model to explain the elongation of vertebrate embryo [Regev et al 2022]. Till now we have described the dynamics of the cells in the PSM in general. Now in the following subsections we will consider specific cases to understand the dynamics and mechanical properties of the PSM.

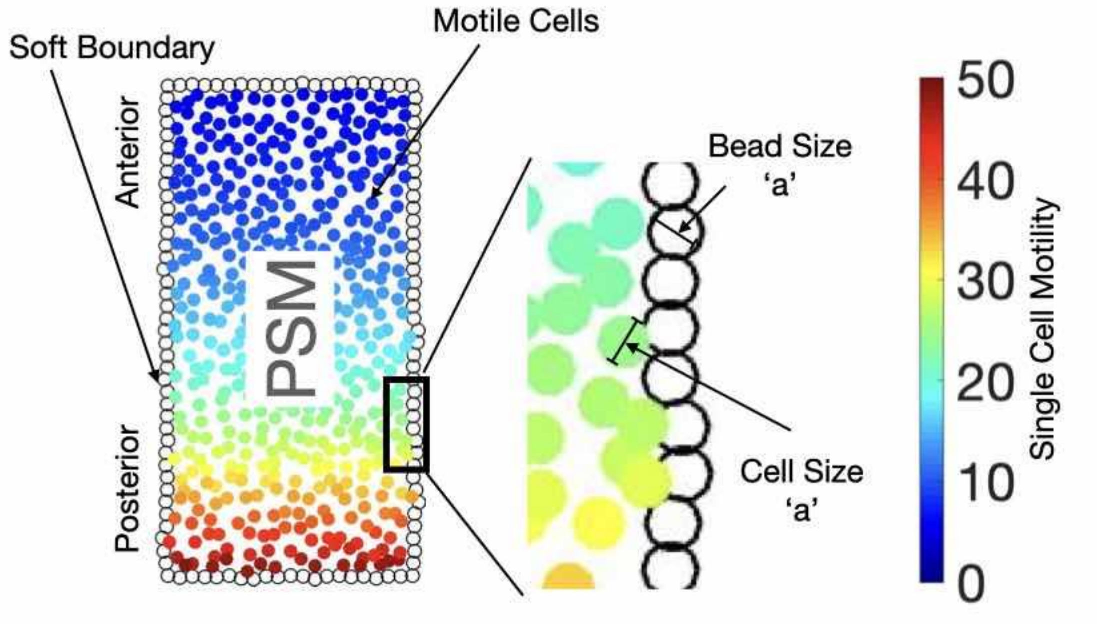


Fig. 1: 2D Model for the ex vivo PSM dynamics. We can control the profile of the single cell motility and the absolute value of single cell motility to control the dynamics of the PSM.

**a> PSM *ex vivo*:**

We consider that the PSM is confined in a soft boundary that is made up of the beads of the same size as the cell, *i.e.*, $a$ and these beads are connected to each other with a spring with spring constant k = 50 Fig 1 (b). The interaction among the beads of the soft boundary and the cells of the PSM is purely repulsive in nature and the force acting on the single cell with position coordinate $\boldsymbol{r}_i$ can be given by

$$f_{ext}^{i} = \sum_{j} \begin{cases} -k(\boldsymbol{r}_i - \boldsymbol{r}_j - a) & \text{for } r_{ij} \leq a \\ 0 & \text{for } r_{ij} > a \end{cases}$$

Where $\boldsymbol{r}_j$ is the position of the $j^{th}$ bead and $r_{ij}$ is the separation between cell $i$ and bead $j$. We use the overdamped Langevin equation to describe the dynamics of the beads without the noise term,

similar to the equation to describe the dynamics of the cell with a different intra-species (bead-bead) interaction.
This setup is to mimic the *ex vivo* dynamics of dissected PSM. We start the simulation with an aspect ratio of 19 and with 1020 cells in the PSM and 400 beads in the soft boundary. We do not consider any cell division while we are monitoring the dynamics of the PSM, so the number of cells in the PSM remains fixed. The single cell motility value close to TB is $\mathcal{M} = 50$ and then the motility values decay exponentially with the position of the cells away from the TB with the length scale $L = a\rho_0\mathcal{D}\tau$. All the length scales are rescaled with the cell size, so we choose $a = 1$. We choose $\rho_0 = 0.65$, $\mathcal{D} = 1.0$ and $\tau = 50$. The initial shape of the PSM is a rectangle with aspect ratio 19, but with time the PSM starts to shrink in total length and its width increases in the posterior end (Fig. 2B). We have shown in Fig. 1B that tissue adopts a pear-like shape, similar to what we have observed in the experiments. The colorbar shows the single cell motility $\mathcal{M}$ of the cells in PSM.

To get a better handle on the mechanical propertiy of the PSM along AP direction we performed simulations by breaking the PSM into three parts, the posterior, i.e., highly active one (pPSM), the mid region, i.e., moderately active (mPSM) and the anterior, i.e., least active one (aPSM). For all three cases, we have chosen the aspect ratio to be 2, and this parts of PSMs are confined with a rectangular ($40a \times 20a$) soft boundary before the simulation begins. In these simulations we used the same parameter as we have used for the dynamics of the full PSM. We see that these parts of PSM starts to become circular with time due to fluidity of the tissue and the effective surface tension of the soft boundary. Since the fluidity of all the three parts are different, the time scale to relax to the circular shape is also different (Fig. 2, C and D). The pPSM relaxes fastest and aPSM relaxes slowest. We have also shown the snapshots of the evolution of the parts of PSM at different time points (Fig. 2C). Next we looked at the mean squares displacement (MSD) for all the three cases and using this data we extracted the diffusivity $D$ in these three regions using the relation:

$$< \Delta r^2(\Delta t) > = 4D\Delta t + \mathrm{v}^2\Delta t^2.$$

Once we have diffusivity $D$, we used the Stokes-Einstein equation which is valid in low Reynold numbers to extract the viscosity $\eta = \frac{k_B T}{6\pi D a}$. Where $k_B T$ can be replaced by the average activity of the region $\mathcal{M}$. In Fig. 2D, we show the viscosity in all the three regions and for posterior region the viscosity is least, whereas for anterior region the viscosity is maximum. These results are in qualitative agreement with our experiments.

### b> PSM in a Channel:

In this case we consider that the PSM is confined in a microchannel with a rigid wall. The walls of the channel have reflecting ($\boldsymbol{f}_{ext}$) boundary condition (Fig 2E). To begin with we start with a channel of width $10a$ and PSM is confined in this channel, the posterior end of the channel is completely sealed, so if there is going to be any growth or shrinkage to the PSM it will be observed from the anterior end. We are using the same parameter for the PSM as we have used for the PSM which was confined within a soft boundary. For this case we have 507 cells in the PSM. The evolution of the PSM in a channel is shown in Fig. 2B at different time points and we observe that first PSM shrinks a little and then PSM grows with time, which is in qualitative agreement with the experiments. To compare this growth for a different channel width ($20a$) we took the same PSM and put it in the microchannel, since we are using the same PSM and doubling the channel width which means that the decay length scale will become half now. When we compared the growth of the PSM for these two channel widths we found that the PSM in narrower channel grows faster than the wider channel (Fig 2F). In Fig. 2C, we have reported $\Delta l = l - l_{min}$ and $\Delta t = t - t_{min}$, where $l_{min}$ is the length of the PSM when it is minimum and $t_{min}$ corresponds to the time when $l_{min}$ is observed.

Then we measured the stress applied by these PSMs when it evolves in a channel and in an embryo. To mimic the dynamics of PSM in an embryo, we must also consider the fact that there is addition of new cells from the tailbud (TB). We have used the same approach as described in Regev *et.al*. 2022, and to mimic the addition on the cells from TB, we use the condition that as soon as there is a gap between the TB and the PSM that is large enough to accommodate a new cell of size $a$ we add a cell to the PSM. To measure the stress, in the beginning we put a wall that is attached to a spring and is pressing against the PSM at the posterior end in the AP direction. As PSM grows it will start to apply a force against the wall and since the wall is attached with a spring the resistance against the force applied by the PSM will increase with the growth of PSM and eventually there will be an equilibrium between the resistance force of the wall due to the attached spring and the forces which is applied by the embryo and that will give us the point of maximum stress applied by the PSM. We observe that the stress applied by the PSM for the embryo is larger than the that of the channel (Fig. 2G-H) and this increase in the stress is attributed to the addition of new cell.

We performed another simulation with reduced motility to understand the effect of inhibition of FGF. To do this we reduced the single cell motility $\mathcal{M}$ by a factor of 2.5 and we observe that the growth of PSM with reduced motility decreases significantly (Fig. 2F), which can be attributed to the fact that the force which is required to grow comes from the active pressure and since $\mathcal{M}$ decreases which implies that the active pressure decreases and hence the growth of the PSM. We also measured the stress applied by the PSM for the reduced motility case and we find that it reduces significantly (Fig. 2H). These results are in qualitative agreement with our experiments.
